# Circuits activated by psychiatric-associated behavior: from brain-wide labeling to regional assessment using Psych-TRAP

**DOI:** 10.1101/2025.08.24.671963

**Authors:** Meet Jariwala, Xhuliana Sula, Emma Kathrine Hjarding, Sofie Braum Holst, Konstantin Khodosevich

**Affiliations:** Biotech Research and Innovation Center, BRIC, University of Copenhagen, 2200 Copenhagen, Denmark

## Abstract

Genetic risk factors are major contributors to psychiatric disorders, yet limited knowledge exists about how these risk factors impair brain circuits to produce disease phenotypes. Better understanding of this process will help us identify potential drug targets acting directly on the vulnerable cell types and circuits. To elucidate how risk factors impact brain circuitry in relation to psychiatric phenotypes, we employed a clinically relevant mouse model of a severe psychiatric genetic risk factors - the 15q13.3 microdeletion. We developed a novel technique, Psych-TRAP, to label psychiatric behaviour-associated circuits at whole brain level and with cellular resolution. Using Psych-TRAP, we permanently labelled transiently active cells responding to a psychiatric-relevant behavior (social interaction) and quantified their densities across the whole brain. We validated Psych-TRAP by confirming activation in previously known brain areas involved in social interaction. Furthermore, we discovered that impaired social interaction behavior in 15q13.3 microdeletion mice was associated with alterations in the GABAergic component in the activated circuit, mainly mediated by reelin-positive neurons in the prefrontal cortex and by somatostatin-positive neurons in the somatosensory cortex. Thus, Psych-TRAP represents a robust technique to permanently label transiently active cells during behavior in psychiatric risk factor models for future circuit manipulations in an unbiased manner that will help to reveal the underlying molecular markers mediating psychiatric behaviours.

## Introduction

Psychiatric disorders comprise a broad group of disorders with a variety of complex behavioral manifestations. [1][2]. These manifestations involve problems with emotional regulation, learning, memory, motor skills and others [1]. Each of these behavioral impairments are regulated by specific circuitry, both at local and long-range levels. Elucidation of these circuitries is crucial to study the types of neurons that govern specific behaviours and design therapeutics that can restore behavioral phenotypes [3].

The brain has highly complex structure, consisting of thousands of neuronal subtypes located in brain regions that often have specialized functions. Indeed, recent studies show the immense complexity of neurons that is established over course of brain development and maturation when most psychiatric disorders arise [4][5][6]. Due to such complexity, it is hard to understand exactly what circuits and neurons within them contribute to behavioral outcomes, and the challenge of studying psychiatric phenotype adds another layer of complexity. A promising strategy is to map assemblies of neurons that are activated during a specific behavior and characterize these mapped neurons at whole brain level. Comparing mapped neurons in wild-type mice and those that model specific psychiatric risk factors can help to identify disrupted circuits and brain areas and explain behavioral deficits related to psychiatry-associated phenotypes. Currently, most common methods to label activated neuronal assemblies with a strong spatial-temporal resolution rely on expression of immediate early genes (IEGs) [7][8][9], Ca2+ concentration dependency [10][11] or release of synaptic vesicles [12][13] as a measure to indicate neuronal activity. However, no such technique, to date, has been implemented to label neurons that are responsible for behavioral impairments at whole brain level and in a face-validated genetic model of psychiatric risk factor.

In this study, we propose a novel paradigm (Psych-TRAP) to study psychiatric-associated behaviours and to permanently label neurons activated during behavior throughout the whole brain with high spatial-temporal resolution. A recently developed technique, TRAP2 [7], captures transiently expressed IEG, namely *Fos*, making it possible to label active neurons and circuits permanently [14]. Combining TRAP2 with any robust face-validated psychiatric mouse model will help to dissect behavioral symptoms at circuit-level. To this end, we implemented TRAP2 technique to 15q13.3 microdeletion model mice that proved to be a robust neuropsychiatric model [15][16]. We captured neurons using TRAP2 and applied whole brain imaging and region-specific quantification of activated neurons using ABBA [17], an automated tool for brain area registration and labelled neuron quantification to validate the activation of neurons in specific brain areas. Further, we validated neuron-type specific activation patterns that underlie behavioral impairments. Therefore, we offer here a rationale for researchers in psychiatry field to study circuits at single neuron resolution level that coordinate behavioral impairments associated with psychiatric disorders.

## Methods

### Animal husbandry

Experimental procedures on animals were conducted in accordance with Danish legislation and guidelines published by the National Animal Ethics Committee of Denmark. The TRAP-WT mice were generated by crossing TRAP2 mice (Jackson Laboratory: 030323) with Ai9 reporter mice (Jackson Laboratory 007909). To create the triple transgenic animal line, the TRAP-WT mice were crossed with (Df(15q13)/+) mice (Taconic Biosciences 10962) yielding TRAP-15q^+/-^ mice. All genes were kept in heterozygous state and animals containing homozygous genotypes were not included in the study. The TRAP-WT animals in this project were TRAP2; Ai9 mice that did not contain the 15q13.3 microdeletion. Mice were housed together in individually ventilated cages with ad-libitum access to food and water along with sawdust and other nesting materials in all cages. The conditions (temperature regulation, ventilation and humidity) were constantly controlled in a reversed 12-hour cycle. The mice used for the experiment were older than 60 days, which is equivalent to minimum 20 years in humans [5].

### Genotyping

Mouse genotypes were determined by polymerase chain reaction (PCR) using Accustart II Polymerase (QuantaBio). PCR conditions were as follows: 94 °C for 2 min, followed by 35 cycles of 94 °C for 15 s, 62 °C for 30 s, 72 °C for 30 s, and final extension at 72°C for 5 min. The primers (in 5’-3’) (TAG Copenhagen A/S, Denmark) used for detection were as follows:

TRAP2 wild forward: GTC CGG TTC CTT CTA TGC AG (357 bp)

TRAP2 common: GAA CCT TCG AGG GAA GAC G (heterozygote 357 bp + 232 bp)

TRAP2 mutant forward: CCT TGC AAA AGT ATT ACA TCA CG (232 bp)

tdTomato (Ai9) wild type pair (AAG GGA GCT GCA GTG GAG TA and CCG AAA ATC TGT GGG AAG TC)

tdTomato (Ai9) mutant pair (GGC ATT AAA GCA GCG TAT CC and CTG TTC CTG TAC GGC ATG G)

Df(15q13)/+ wild type pair (CTGTATCCGTAAACACGCTAACTGG and GAGCAGTGTTTGTGTGGTTTATGG)

Df(15q13)/+ mutant pair (TCGAGCCTAGGCATAACTTCGTATA and TCGGCACCTTGATGAGTCTTG)

### Solutions and Drugs

4-Hydroxy tamoxifen (4-OHT) (H-6278, Sigma-Aldrich) solution was dissolved in dimethyl sulfoxide (DMSO) (154938, Sigma-Aldrich) to create stock solutions and was kept at -20℃. Fresh 4-OHT solution was prepared under a chemical hood by thawing 4-OHT-DMSO solution, mixed with 2% Tween80 (Sigma-Aldrich P1754), diluted in 0.9% saline and always protected from light. Animals were injected intraperitoneally according to weight with 30 mg/kg of an adult mouse. Because of the harmfulness of 4-OHT, being carcinogenic in nature, all procedures were carried out while wearing adequate protective equipment and the waste was discarded according to animal authority guidelines given by the Department of Experimental Medicine, Copenhagen University.

### Behavior

Mice were habituated for at least four days prior to the actual experimental day to avoid unnecessary stress to the animal. This was done by single housing the test mice to ensure a strong need for social interaction and to avoid stress and injuries due to fighting among littermates. Test animals were habituated for being lifted and pinched gently on the lower abdomen to mimic the 4-OHT injection process on the test day, as sudden stress on the test day could influence activation of TRAP+ neurons. Stranger mice were also habituated for 4-7 days for 10 minutes to sit calmly inside the metal cage. A 3-chamber social interaction (3-CSI test) was used to assess the mice’s social interaction as shown in **Fig 1C**. 3-CSI consists of three chambers, and each chamber is separated by a glass wall with a small opening enough to allow free movement of mice. The stranger mouse was always chosen as the younger mouse of the same gender to avoid conflicts due to social hierarchy. The test mouse was placed in the buffer zone, and the behavior of the mouse was then tracked for 10 minutes. Between every behavior, the apparatus was cleaned using 70% ethanol and wiped with dry tissue paper. Animals were injected with 4-OHT solution (30mg/kg) right after the 3-CSI test and were housed in disposable cages for 7 days undisturbed.

**Figure 1:**
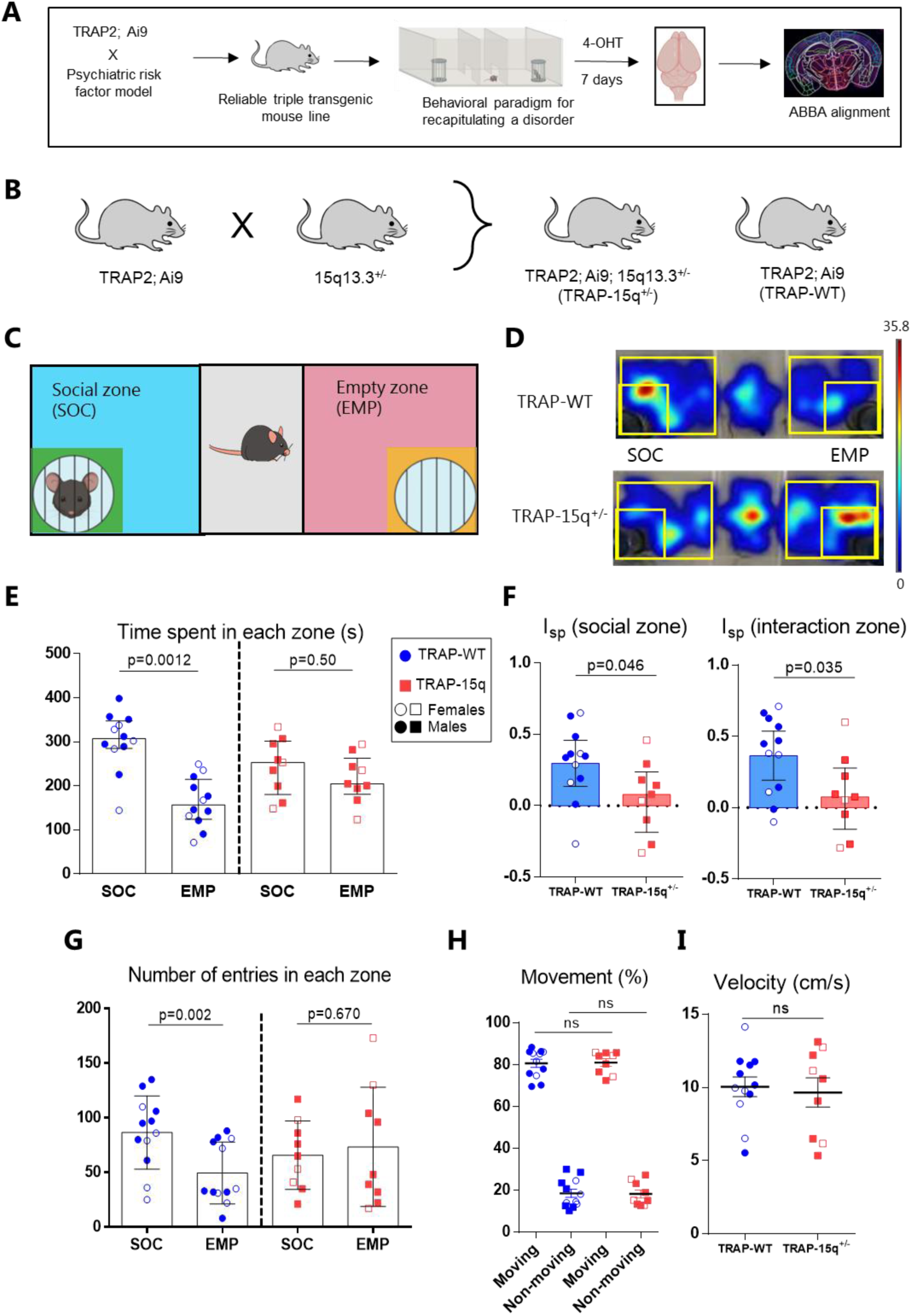
The Psych-TRAP technique. A: An overview of the Psych-TRAP technique, including steps to breed an intersectional transgenic mouse line, label active cells, align brain sections and quantify in an automated manner. B: Generation of a triple transgenic mouse line, yielding to TRAP-15q^+/-^ and TRAP-WT littermates. C: 3-chambered social interaction test chamber (3-CSI). Blue and pink chambers are social (SOC) zones and empty zones (EMP). Green and yellow zones are their respective interaction zones. D: Heatmaps of two representative animals from TRAP-WT and TRAP-15q^+/-^ groups showing the amount of time spent in the chamber. Color bar represents red as high intensity and blue as low intensity. E: Time spent by animals in each zone in seconds (s). Blue circles and red squares indicate TRAP-WT mice and TRAP-15q^+/-^ mice, respectively. Male and female mice are denoted by filled and clear circles/squares, respectively. SOC: social zone, EMP: empty zone. (TRAP-WT: n=12 mice, SOC: 302.1±19.08, EMP: 162.6±16.16, paired t-test p=0.0012; TRAP-15q^+/-^: n=9 animals, SOC: 243.8±21.43, EMP: 216.4±18.00, paired t-test p=0.50). F: Social preference index (I_sp_) of social zone (TRAP-WT: n=12 mice, 0.297±0.074; TRAP-15q^+/-^: n=9 mice, 0.055±0.085, unpaired t-test with Welch’s correction p=0.046) and interaction zone (TRAP-WT: n=12 mice, 0.365±0.08; TRAP-15q^+/-^: n=9 mice, 0.088±0.092, unpaired t-test with Welch’s correction p=0.035). G: Number of entries in each zone (TRAP-WT: n=12, SOC 86.58±9.68, EMP 49.50±8.15, Wilcoxon test p=0.002; TRAP-15q^+/-^: n=9, SOC 65.78±10.48, EMP 73.44±18.23, paired t test p=0.670). H: Movement (%). Unpaired t-test with Welch’s correction. I: Velocity (cm/s). Unpaired t-test with Welch’s correction. All data are represented as ± S.E.M, p < 0.05.

### Behavior analysis

Behavior was recorded with a ceiling-mounted camera and later scored by the EthoVision XT software (Noldus). The software was trained to track the nose point of the test mice. Automated tracking was followed by a manual review of the footage by the experimenter, to ensure that software-derived mistakes are corrected. Active interaction was considered when the mouse directly interacts in proximity to metal wired-cage, directly faces towards it and sniffs around the wired-cage, otherwise all activity was considered as non-interaction. The experimenter and behavior scorer is blinded to the genotype during the behavior and scoring.

### Brain fixation

Mouse brains were collected after 7-10 days of 4-OHT injection for histological preparations. Mouse brains were fixed for immunohistochemistry first by anesthetizing using Ketamine (100mg/ml)-Xylazine (20mg/ml) cocktail (Department of Experimental Medicine, Copenhagen University), then by performing transcardial perfusion with cold phosphate-buffered saline (PBS) to flush out blood and at last with 4% paraformaldehyde (PFA, Roth) to fix the brain. Brains were removed, postfixed in 4% PFA overnight and were transferred to PBS containing 0.02% sodium azide (Sigma S2002) at 4℃ until further processing.

### Vibratome sectioning and immunohistochemistry

Collected brains were sliced using a vibrating microtome (Leica VT1000S). Brains were sliced free-floating with 100 microns thick coronal sections and were stored in PBS with sodium azide at 4℃. Immunohistochemistry was performed by washing selected sections in PBS briefly and incubating in blocking buffer (10% normal donkey/horse serum (Gibco 16050-122), 0.2% Triton X-100 (VWR International, Denmark) in PBS) at room temperature for 2 hours on a shaker. Subsequently, sections were incubated overnight at 4℃ with primary antibodies made in another buffer (2% normal donkey/horse serum, 0.2% Triton X-100 in PBS) on a shaker. The following day, sections were washed in PBS and incubated with Alexa Fluor conjugated antibodies (Invitrogen) (in the same buffer as primary antibodies were incubated with) either for 2 hours at room temperature or 24 hours at 4℃ on a shaker. Nuclei were counterstained with DAPI (1:50000, Sigma) for 10 minutes and slices were mounted on Superfrost microscopic slides using FluoroMount-G medium (ThermoFischer Scientific) or Immu-mount (Epredia). After mounting, the slides were stored in a 4℃ fridge in a closed container to protect it from light. Primary antibodies used here: anti-somatostatin (mouse, 1:1000, Santa Cruz sc74556 and rat, 1:500, Millipore MAB354), anti-parvalbumin (goat, 1:2000, Swant PVG213 and mouse, 1:500, Sigma P3088), anti-reelin (goat, 1:500, R&D Systems AF3820), anti-GAD67 (mouse, 1:500, Millipore MAB5406), anti-Iba1 (rabbit, 1:250, Wako 019-19741), anti-GFAP (mouse, 1:2000, Sigma G3893), anti-NeuN (mouse, 1:1000, Millipore MAB377 and chicken, 1:1000, Millipore ABN91).

### Imaging

Imaging was done using Zeiss confocal microscope LSM800. The 25x objective with 0.5x magnification was used for all images. Z-axis was set for maximum intensity of signals. For ABBA slices, 10x objective was used to capture the whole slice and was made sure that all parts of slice are in focus. Consistent parameters were maintained across all images. All Images were analysed in Fiji [18].

### ABBA alignment and quantification

Sections were washed in PBS, counterstained with DAPI and mounted on microscopic slides. The pipeline was followed as mentioned in the original article [17]. Approximately 16-19 sections per brain, with approximately 600 microns between two corresponding slices, were picked anterior to posterior (AP: +3.0 mm to -5.0 mm) to get an overview of the whole brain. The sections were added to a project file in QuPath and imported to ImageJ Fiji using ABBA plugin. Successful alignments were performed in ImageJ Fiji using different registration strategies (Affline and Spline) to match the edges of brain sections and sub-areas based on Nissl staining provided in the plugin. Once aligned, the images were exported to QuPath and were used for detection of for positively labelled cells. Cell detection parameters for ABBA were optimized to get consistent positive cells across the sections and across all animals. Exported files contain number of detected cells, area and labels of each brain area. Python scripts were used to normalize to the area generating heatmaps.

### Statistical analysis

Statistical graphs were generated in GraphPad Prism 6 and Python. All statistical tests were performed using Python scripts. All data were tested for normality using Shapiro-Wilk normality test. Appropriate post hoc tests were carried out for paired and unpaired data. Heatmaps for ABBA whole brain quantification was generated by normalizing the raw cell counts per slice with area of the brain region. Brain areas that were missing in at least one animal (due to fracture of brain sections, break during sectioning/taking brain out of the skull, or folding of slices), were excluded to have an unbiased cell count quantification. Normalized cell densities were transformed into log2 (x+1), where x is cell density of each brain area. Medians of each group were compared by Mann-Whitney U test. We did not perform False Discovery Rate (FDR) analyses where we had less than 6 mice per group. We did not perform power calculations to estimate the size of our animals per group. Female and male mice were analysed together and were ensured that all animals came from at least 3 independent litters. Animals were excluded based on the following criteria: died during habituation or behavior (n=1), died post injection of 4-OHT (n=1), leakage of 4-OHT while injection (n=4), error video file (n=1), inefficient perfusion or difficult to image (n=3). Complete statistics are provided in **Supplementary File 1**.

## Results

### Framework to label psychiatric-associated circuits

Here, we propose a combinatorial technique, Psych-TRAP, to tag transiently activated neurons followed by clinically relevant behavior in a mouse psychiatric risk factor model, and to align with the available structural brain atlas [19] in a brain-wide single neuron resolution manner for automated alignment and annotation. We implemented this technique to study brain region-specific changes in neuronal activation that are associated with behavioral impairments (**Fig 1A**). We employed this approach for an intersectional triple transgenic mouse line that reliably recapitulates clinical behavioral symptoms and is easy to breed and maintain. To this end, we crossed the TRAP2; Ai9 reporter mice with mouse model of 15q13.3 microdeletion syndrome (**Fig 1B**). The 15q13.3 locus have been associated with severe neuropsychiatric brain dysfunctions in humans and has complex genetic architecture, high penetrance, gene-dosage dependent manifestations of phenotypes [20][21][22]. Mouse model for 15q13.3 microdeletion (Df(h15q13)/+ recapitulates well behavioral and functional changes observed in the patients, both in 15q13.3 microdeletion syndrome and in general in neuropsychiatry [23][24][15][16]. We refer to this intersectional transgenic model as TRAP-15q^+/-^ (**Fig 1B**).

To visualize the cells activated after a specific stimulus or behavior, we employed TRAP2 mouse model [7] crossed with a tdTomato reporter background (TRAP2; Ai9). TRAP2 uses a drug-mediated approach (4-Hydroxy Tamoxifen, 4-OHT) to target inducible CRE (iCre-ER^T2^) under the FOS promoter, thereby activating the *c-fos*, an immediate early gene, which in-turn activates the floxed tdTomato protein by targeted recombination, exclusively in the active cells [7]. Such approach leads to permanent labeling of activated cells.

### Behavioral signatures of TRAP-15q^+/-^ mice recapitulate disrupted social interaction behavior

To deeply understand the behavioral aspects and validate clinically relevant behavioral symptoms, we subjected the mice carrying TRAP-15q^+/-^ microdeletion and mice without the microdeletion (TRAP2 wild type [TRAP-WT] littermates) to a 3-chambered social interaction (3-CSI) paradigm [25]. The chamber consists of three zones - social zone (SOC), buffer zone and empty zone (EMP) (**Fig 1C**). SOC harbors a younger mouse of the same sex inside a cage with metal bars to initiate an active social interaction, whereas EMP harbors the empty cage.

As expected, TRAP-WT mice spent relatively more time in the SOC as compared to EMP, indicating a normal sociability. On the contrary, TRAP-15q^+/-^ animals showed no such preference in time spent between SOC and EMP (**Fig 1D, E**), indicating a disruptive social interaction behavior. We further zoomed into the interaction zone (green and yellow color zones around the cage) of the 3-CSI test (**Fig 1C**) and quantified the differences in the time spent by the animal while performing active interaction around the metal cages using a social interaction index (I_sp_). Again, we observed that TRAP-WT animals spent significantly more time in active interaction, whereas TRAP-15q^+/-^ mice showed little or negative preference (**Fig 1F**).

We also quantified how frequent an animal enters the interaction zone in SOC and EMP. We found that TRAP-WT animals made significantly higher numbers of entries in the SOC than in EMP, while TRAP-15q^+/-^ animals showed no preference, indicating a lack of motivation to interact actively (**Fig 1G**). Lastly, to exclude influence of movement impairments on the observed phenotype in the 3-CSI test, we evaluated movement and velocity of both animal groups and observed no differences (**Fig 1H, I**), confirming that the lack of social preference is exclusively due to the 15q13.3 microdeletion. To determine potential sex-specific effects of 15q13.3 microdeletion, we separated females and males for behavioral analyses and found no obvious effects in any behavioral parameters (**Fig 1E, F**).

Overall, we have successfully demonstrated that TRAP-15q^+/-^ transgenic model can be reliably used for our strategy to label and investigate circuitry that is activated by psychiatric-associated behavior.

### Whole-brain automated registration and quantification of activated neurons

To identify circuitry that was activated during social interaction behavior at a brain-wide scale, we aimed to find an accurate and efficient technique for automated brain sections alignment with available brain atlases allowing for whole-brain quantification with regional single neuron resolution. We injected 4-OHT immediately after the 3-CSI behavior and fixed the brains 7 days postinjection to allow targeted recombination in the active cells. We aligned approximately 17-19 coronal brain sections per mouse from the whole brain with the available Allen Mouse Brain Atlas [19] (**Fig 2A**). These sections were ensured to be corresponding sections from anterior to posterior regions in all animals to maintain the consistency. Representative brain sections (5 sections) are shown in **Fig 2B and Fig S1**, indicating successful alignments of our imaged slices (left) with the mouse brain atlas in anterior to posterior direction (right).

**Figure 2:**
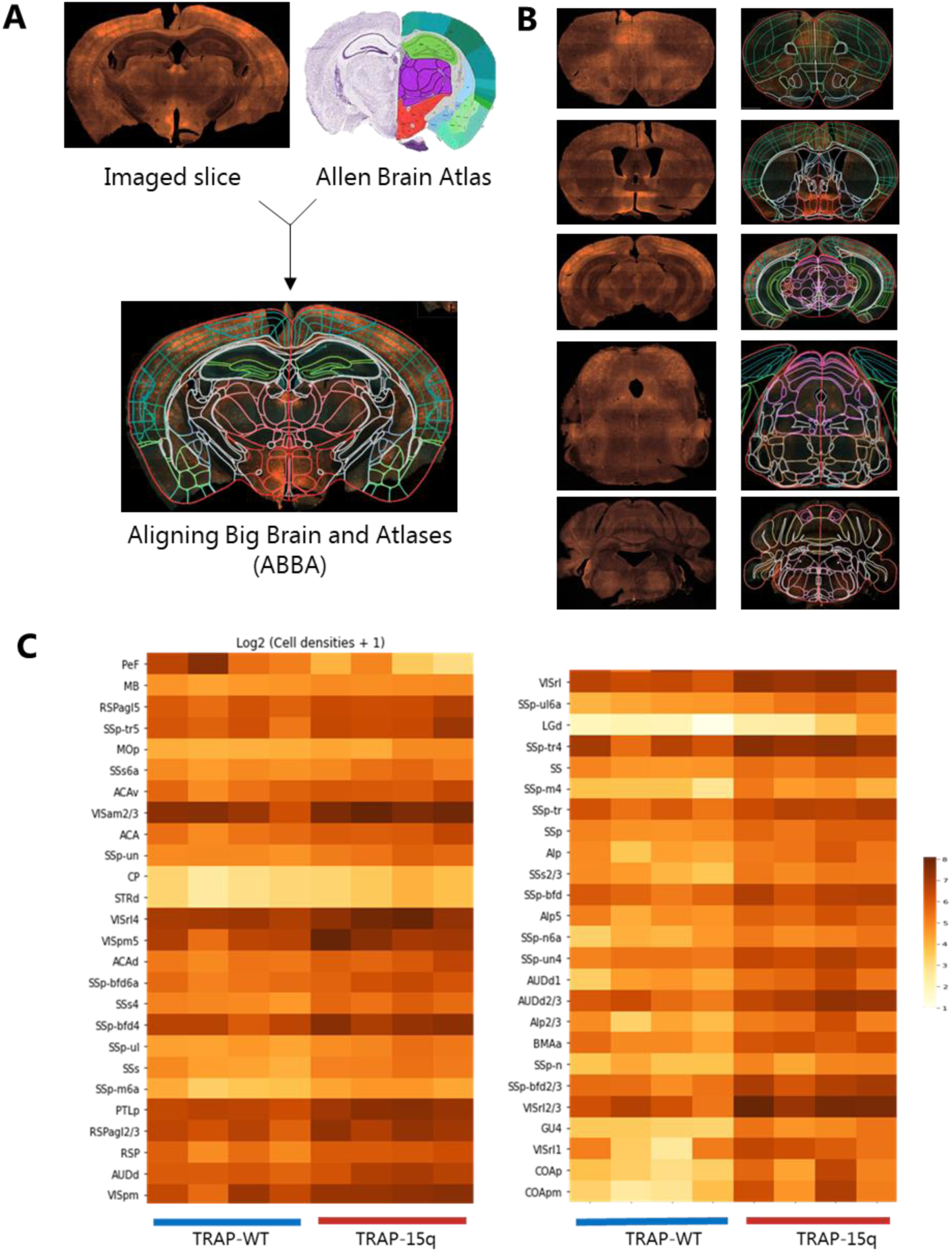
Automated alignment, registration and quantification using ABBA. A: Strategy to combine imaged brain section (left) and corresponding section from Allen Brain Atlas (right), yielding an example of a successful alignment using ABBA (below). B: Representative examples from 5 brain sections, anterior to posterior (top to bottom), confirming successful alignments using ABBA. C: Heatmaps of log2(cell densities +1) showing 52 brain areas from TRAP-WT and TRAP-15q^+/-^ mice (4 mice per group, blue and red lines below indicate TRAP-WT and TRAP-15q^+/-^, respectively) that showed a differential activation of cells, Mann-Whitney U test, p < 0.05.

With the help of ABBA (<u>A</u>ligning <u>B</u>ig <u>B</u>rains and <u>A</u>tlases), an ImageJ FIJI plugin [17], we quantified in an automated manner the active cells, as observed by tdTomato expression (**Fig S1**), from all the successfully aligned brain sections. After optimization of alignment procedure and filtering out the broken section/regions, we identified 474 brain regions that could be used for the analysis of activation pattern of cell densities normalized to the area of measurement. The heatmaps in **Fig 2C** (raw counts in **Fig S2**) show top 52 brain regions where we found a systematic difference in the density of activated cells after 3-CSI between TRAP-WT and TRAP-15q^+/-^ mice.

We observed several cortical areas, including specific layers, showing a higher trend of activated cells (**Fig 2C**). The brain regions that showed highest differential activation of cells after social interaction behavior were somatosensory areas, both in primary somatosensory areas (SSp) and supplemental somatosensory areas (SSs) (subregions: SSp-tr5, SSs6a, SSp-bfd6a, SSp-bfd4, SSp-ul, SSp-m6a, SSp-ul6a, SSp-tr4, SSp-m4, SSp-tr, SSp, SSs2/3, SSp-bfd, SSp-n6a, SSp-un4, SSp-n, SSp-bfd2/3). Other cortical areas that exhibited high differential activation of neurons in TRAP-15q^+/-^ animals as compared to TRAP-WT animals were visual areas (VISam2/3, VISrl4, VISpm5, VISpm, VISrl, VISrl2/3, VISrl1), retrosplenial area (RSP) and its sub-regions (RSPagl5, RSPagl2/3), auditory areas (AUDd, AUDd1, AUDd2/3), one sub-region of gustatory area (GU4), anterior cingulate area (ACA, ACAv, ACAd), posterior agranular insular areas (AIp, AIp5), primary motor area (MOp), posterior parietal association area (PTLp) and olfactory areas (posterior cortical amygdala areas -COAp, COApm).

Among subcortical areas, we found only anterior basomedial amygdalar area (BMAa) that showed differential activation pattern after 3-CSI in TRAP-15q^+/-^ animals. In the striatal areas, we found higher activation in striatum dorsal region (STRd) which harbors the caudate putamen (CP). Among the brainstem areas, we found higher activation of cells in the whole midbrain (MB), where the dorsal part of the lateral geniculate nucleus (LGd) stands out as the prominent sub-region for higher activation. The only brain region that had opposite activation pattern - higher number of activated cells in TRAP-WT than TRAP-15q^+/-^ animals - was the perifornical nucleus (PeF) that is located deep inside the hypothalamus.

To further validate our findings, we zoomed in on some of the most active and relevant for social behavior cortical regions - primary somatosensory area trunk (SSp-tr), and prefrontal cortex (PFC), harboring anterior cingulate area (ACA), prelimbic area (PL) and infralimbic area (ILA). By comparing the densities of activated cells between TRAP-WT and TRAP-15q^+/-^ animals, we observed significantly higher number of activated cells in the whole SSp-tr of TRAP-15q^+/-^ mice compared to TRAP-WT mice (**Fig 3A, C**). Such trend was preserved across all layers of SSp-tr, raising to the largest change in the deeper layers (L5b and L6) (**Fig 3C**). In the PFC, the overall quantification also showed significantly higher number of activated cells in TRAP-15^+/-^ mice, particularly in the ACA and PL (**Fig 3B, D**).

**Figure 3:**
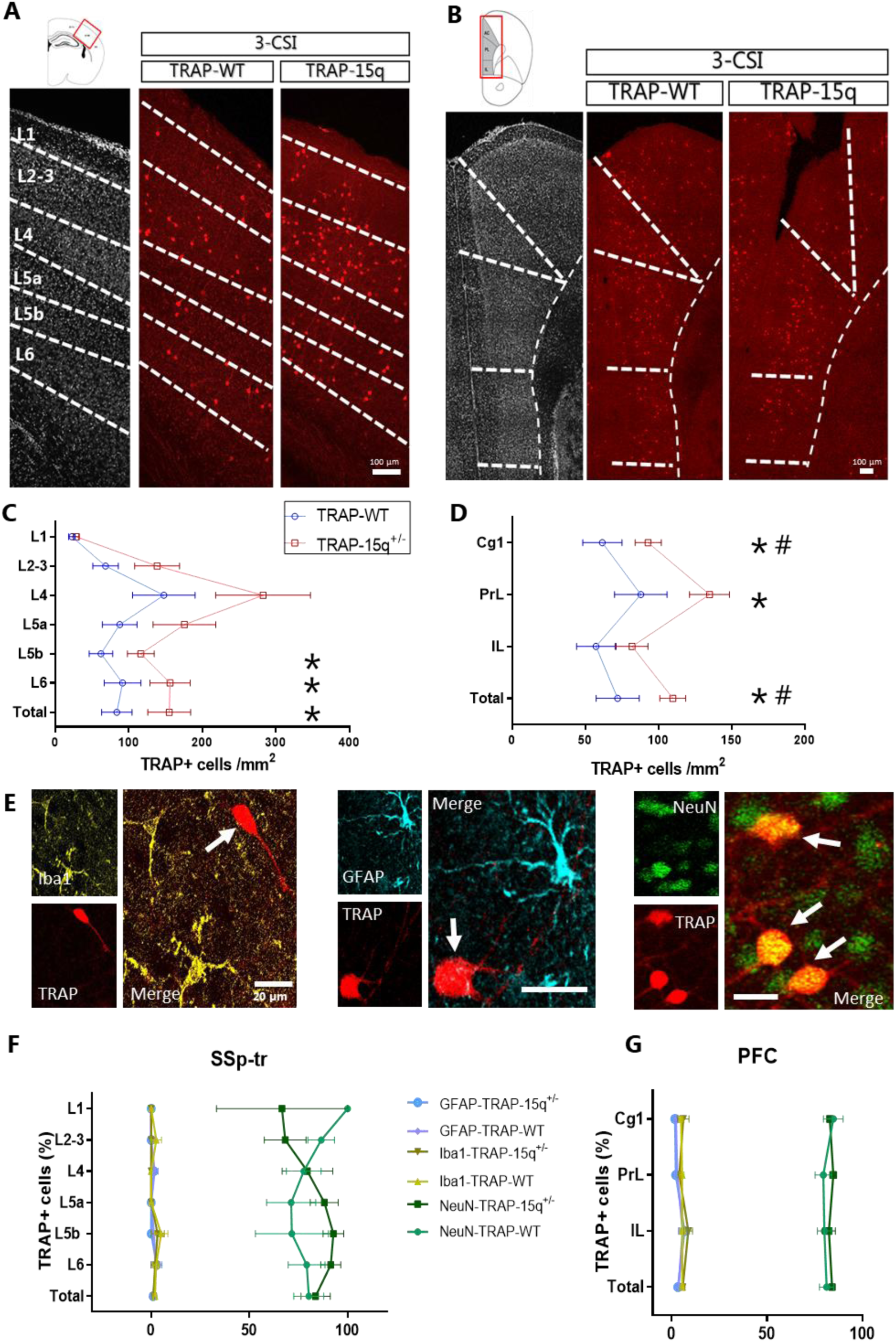
Differential patterns of activated cells in SSp-tr and PFC areas. A: Representative sections of SSp-tr from TRAP-WT and TRAP-15q^+/-^ mice, after 3-CSI and TRAP2-based labeling. Index represents an outline of the brain area where the magnified area is highlighted in red box. First section in gray is DAPI staining to identify cortical layers in SSp-tr. Red fluorescent cells are tdTomato+ cells that are labeled after 3-CSI. Scale bar 100 µm. B: Representative sections of PFC from TRAP-WT and TRAP-15q^+/-^ mice, after 3-CSI and TRAP2-based labeling. Index represents an outline of the brain area where the magnified area is highlighted in red box. First section in grays is DAPI staining to identify cortical layers in PFC. Scale bar 100 µm. C: Comparison of cell densities in layers of SSp-tr between TRAP-WT (n=11) (blue) and TRAP-15q^+/-^ (n=9) (red) mice. Unpaired t-test for all layers, except for L4 (Mann-Whitney U test). D: Comparison of cell densities in sub-regions of PFC between TRAP-WT (n=10) (blue) and TRAP-15q^+/-^ (n=9) (red) mice. Unpaired t-test. E: Representative images of immunostaining against a microglia marker (Iba1, yellow), an astrocyte marker (GFAP, cyan) and a neuronal marker (NeuN, green) along with TRAP+ cells expressing tdTomato. White arrows indicate TRAP+ cells. Scale bar 20 µm. F: Quantification of TRAP+ cells co-expressing Iba1 (TRAP-WT n=5, TRAP-15q^+/-^ n=3), GFAP (TRAP-WT n=3, TRAP-15q^+/-^ n=3) and NeuN (TRAP-WT n=6, TRAP-15q^+/-^ n=4) in SSp-tr in percentages. Mann-Whitney U test. G: Quantification of TRAP+ cells co-expressing Iba1 (TRAP-WT n=5, TRAP-15q^+/-^ n=3), GFAP (TRAP-WT n=2, TRAP-15q^+/-^ n=2) and NeuN (TRAP-WT n=5, TRAP-15q^+/-^ n=4) in PFC in percentages. Mann-Whitney U test. Asterisks (*) indicate p-value <0.05 and Hash sign (#) indicate true FDR <0.05.

Overall, ABBA-mediated automated alignment and quantification provided us with an unbiased overview of several brain areas activated after psychiatric-like behavior. Many of the brain areas from our findings are consistent with those that have been already described as being involved in social behavior [26][27][28][29][30], validating the robustness of this technique. Moreover, we also reported novel brain areas that were not known to play a role in mediating social behavior.

### Characterization of activated cell types

Since tdTomato labeling does not reveal the nature of the activated cells, we characterized the labeled cell types in key regions of interest that displayed major differences in the number of activated cells in TRAP-15q^+/-^ mice the SSp-tr and PFC (**Fig 3C, D**). We first performed labeling of non-neuronal cell markers to determine whether any non-neuronal cells are activated by TRAP2 system in our transgenic mice. To this end, we labeled microglia (Iba1) and astrocytes (GFAP), in the SSp-tr and the PFC. Almost all activated cells were found to be lacking Iba1 and GFAP expression (**Fig 3E-G**) confirming that TRAP2 labeled cells are neurons, which we further validated using the NeuN marker (**Fig 3E-G**). Since inhibitory GABAergic neurons have been shown to make major contributions to brain impairments in psychiatric disorders [31][15][32][33][34], we further focused on determining the subtypes of GABAergic neurons that are differentially activated by social behavior in TRAP-15q^+/-^ mice. We selected three molecular markers for subtypes of GABAergic neurons, namely parvalbumin (PV), somatostatin (SST), and reelin (RELN), owing to previous studies on importance of each of these neuronal subtypes for social behavior and in general altered behavior in psychiatric models [35][36][37][38][16].

Firstly, we studied whether 15q13.3 microdeletion affected distribution of these GABAergic neuronal subtypes in the somatosensory and/or prefrontal cortex. We quantified the cell densities of PV-, SST-, and RELN-positive (PV+, SST+, and RELN+, respectively) neurons in the SSp-tr and overall significant changes were found only in the density of PV+ neurons in the layer 2-3 (L2-3), which was decreased in TRAP-15q^+/-^ mice. Interestingly, there was a visible trend in higher abundance of SST+ neurons in TRAP-15q^+/-^, across all layers, however, it did not reach significance (**Fig 4A, C, E**). In the PFC, there was a striking reduction in the density of PV+ neurons in TRAP-15q^+/-^ mice, which reached above 2-fold difference in the prelimbic area (**Fig 4B, D, F**).

**Figure 4:**
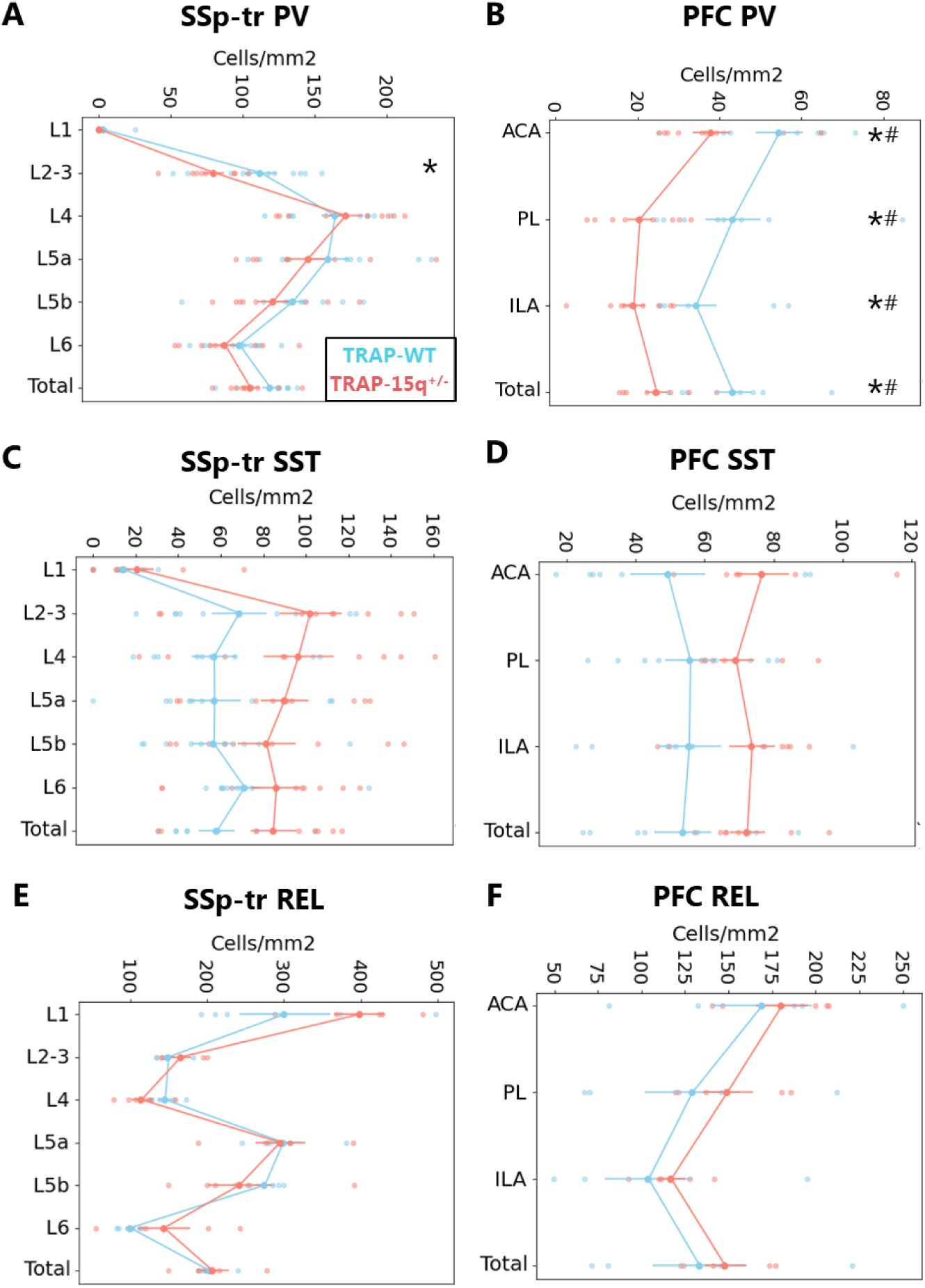
Density of subtypes of GABAergic neurons in SSp-tr and PFC. A-B: Comparison of density of parvalbumin (PV) positive neurons between TRAP-WT and TRAP-15q^+/-^ mice. (TRAP-WT n=10, TRAP-15q^+/-^ n=9), (TRAP-WT n=8, TRAP-15q^+/-^ n=9). Unpaired t-test, except L1 and Total (Mann-Whitney U test). C-D: Comparison of density of somatostatin (SST) positive neurons between TRAP-WT and TRAP-15q^+/-^ mice. (TRAP-WT n=9, TRAP-15q^+/-^ n=9), (TRAP-WT n=8, TRAP-15q^+/-^ n=7). Unpaired t-test, except L1, L6, ACA, PL and ILA (Mann-Whitney U test). E-F: Comparison of density of reelin (RELN) positive neurons between TRAP-WT and TRAP-15q^+/-^ mice. (TRAP-WT n=5, TRAP-15q^+/-^ n=5). Unpaired t-test, except L2/3, Total and ACA (Mann-Whitney U test). Asterisks (*) indicate p-value <0.05 and Hash sign (#) indicate true FDR <0.05. Complete statistics see in **Supplementary File 1**.

Secondly, we investigated whether 15q13.3 microdeletion leads to changes in the activation of GABAergic neurons after 3-CSI behavior. We found a higher density of activated GABAergic neurons in the PFC of TRAP-15q^+/-^ mice, especially in the PL and ILA, suggesting an imbalance in the inhibitory/excitatory system where more GABAergic neurons are recruited in the PFC after 3-CSI behavior (**Fig 5A, C**). Interestingly, we only observed higher trends of activation in SSp-tr of TRAP-15q^+/-^ mice but there were no significant differences in the overall area as well as its sub-regions (**Fig 5B**).

**Figure 5:**
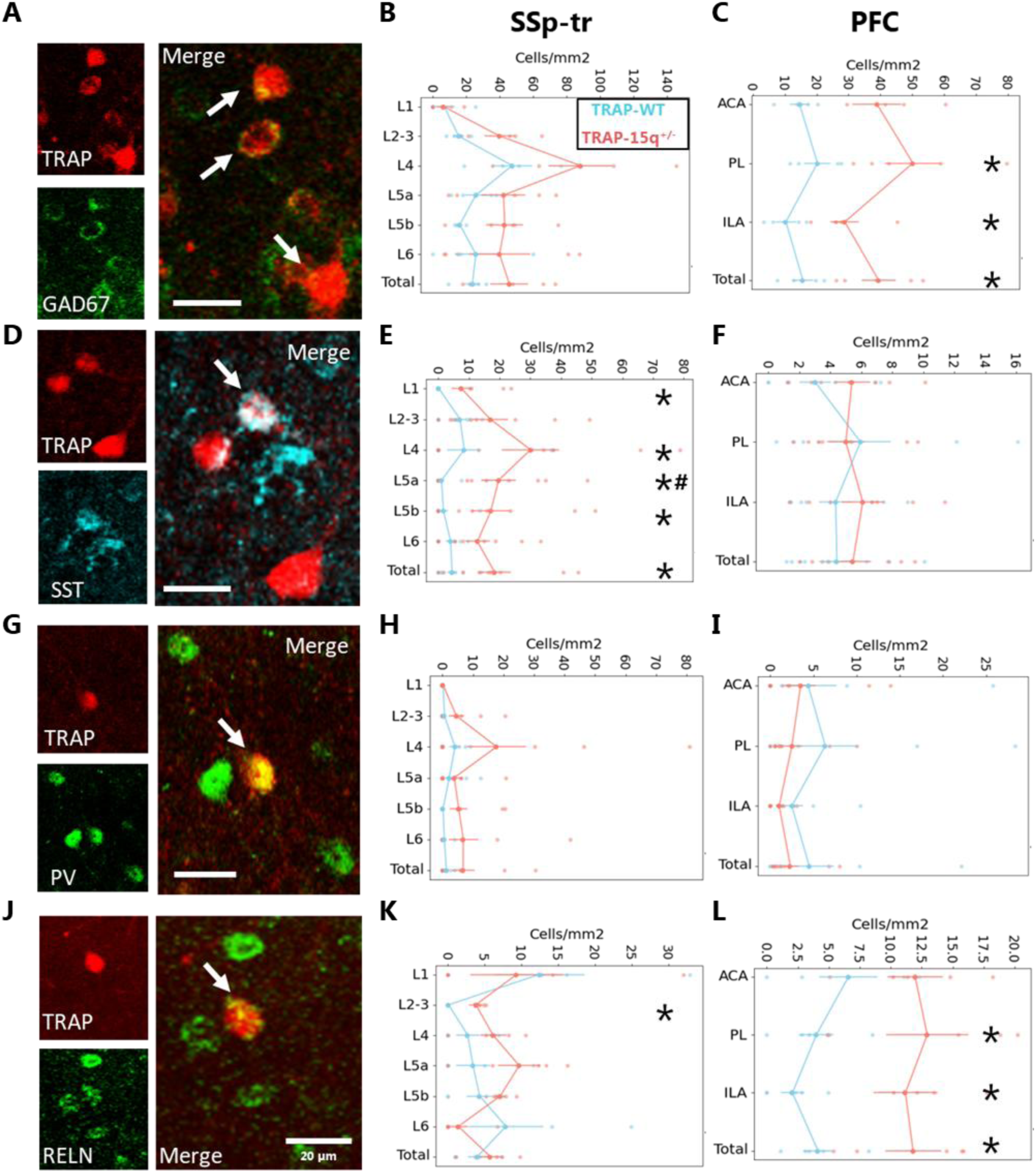
Recruitment of GABAergic neurons in TRAP-WT and TRAP-15q^+/-^ mice after 3-CSI. A, D, G, J: Representative images of immunostaining against GAD67 (A), SST (D), PV (G), and RELN (J) markers. Arrows indicate double positive TRAP+ cells with respective markers. Scale bar 20 µm. B, E, H, K: Quantifications of TRAP+ cell densities co-expressing GAD67 (TRAP-WT n=4, TRAP-15q^+/-^ n=5; Mann-Whitney U test), SST (TRAP-WT n=9, TRAP-15q^+/-^ n=9; Mann-Whitney U test, except L2/3 and Total (unpaired t-test)), PV (TRAP-WT n=10, TRAP-15q^+/-^ n=9; Mann-Whitney U test) and RELN (TRAP-WT n=5, TRAP-15q^+/-^ n=5; Mann-Whitney U test except Total (unpaired t-test)) in SSp-tr. C, F, I, L: Quantifications of TRAP+ cell densities co-expressing GAD67 (TRAP-WT n=4, TRAP-15q^+/-^ n=5; Mann-Whitney U test), SST (TRAP-WT n=8, TRAP-15q^+/-^ n=7; unpaired t-test), PV (TRAP-WT n=8, TRAP-15q^+/-^ n=9; Mann-Whitney U test) and RELN (TRAP-WT n=5, TRAP-15q^+/-^ n=5; unpaired t-test) in PFC. Asterisks (*) indicate p-value <0.05 and Hash sign (#) indicate true FDR <0. Complete statistics see in **Supplementary File 1**.

Lastly, we combined both analyses above and determined whether social behavior in 15q13.3 microdeletion model has specific activation profile of subtypes of GABAergic neurons. We found a selective and consistent pattern of higher density of activated cells co-expressing SST in the SSp-tr in TRAP-15q^+/-^ mice (**Fig 5D, E**), suggesting their crucial role in mediating 3-CSI behavior in the genetic model. Such increase was specific for the SSp-tr, since the PFC showed comparable levels of SST neuron activation (**Fig 5F**). For PV+ neurons, we did not find any changes in activation pattern due to the microdeletion either in the SSp-tr or in the PFC (**Fig 5G, H, I**). Interestingly, while in the SSp-tr RELN+ neurons exhibited only minor changes in their activation pattern (**Fig 5J, K**), in the PFC, there was a dramatic increase in activation of RELN+ neurons, especially in the PL and ILA areas (**Fig 5L**), indicating a potentially important role of RELN+ neurons contributing to impairment in social behavior in 15q13.3 microdeletion model.

Overall, we delineated the differential activation of TRAP+ cells among subtypes of GABAergic neurons and discovered strong changes in activation of SST+ neurons in SSp-tr and RELN+ neurons in PFC in social behavior in the genetic model of psychiatry.

## Discussion

Here, we propose and implement the Psych-TRAP technique to label and quantitatively assess at whole brain level with single neuron resolution, the transiently active neurons that underlie behavioral impairments in a genetic model of psychiatric disorders, (Df(h15q13)/+) [7][24][17]. We have successfully demonstrated potential of Psych-TRAP to determine subtypes of neurons that can be associated with psychiatry-associated behavior. Psych-TRAP technique addresses a major challenge in molecular psychiatry – how to determine what type of neurons and molecular mechanisms within are responsible for specific psychiatric phenotypes at whole brain level. To this end, conventional cell counting in the whole brain is time consuming, requires stereology and thus tedious [39]. Further, Psych-TRAP-labeled neurons can be analyzed downstream by various methods, including classical immunohistochemistry to identify novel targets and brain-wide assessment of vulnerable regions, thus linking behavioral impairments with molecular and cellular mechanisms.

Semi-automated quantification of labeled cells at whole brain level has been optimized by several techniques, such as SHARCQ [40], FASTMAP [41], AMaSiNe [42]. However, each of these techniques also come with limitations of requiring coding and anatomy skills, disproportionate alignment and limited automatic quantifications. In contrast, ABBA [17] offers an automated pipeline to align brain sections automatically using Allen Brain Atlas CCF [19] in an open-source ImageJ Fiji software and integrating the automated quantification into yet another open-source software QuPath.

Psych-TRAP also has promise for connectivity reconstruction at whole brain level. Previous studies used psychiatric models to understand the brain connectivity and activation using f-MRI and BOLD imaging [43][44][45], however this requires expensive instruments and expertise to perform such analyses. Furthermore, resolution is at a regional level. Psych-TRAP provides an unbiased approach to studying any genetic model of psychiatric disorders by TRAP2 technique [7], without compromise on clinical phenotypes, and enables discovery of novel brain areas using ABBA [17], without requiring expertise in coding or anatomy. Potentially, quantitative data can be further utilized to predict connectivity changes. Moreover, additional analyses can “extract” connectivity from the Psych-TRAP-labeled neurons, for instance using single-cell transcriptomics and computational prediction of neuron-neuron connectivity using ligand-receptor interaction [46].

TRAP-WT mice displayed a clear preference towards the stranger mouse, which is expected based on previous studies [47][48][49]. Contrary to TRAP-WT mice, TRAP-15q^+/-^ mice displayed a lack of preference of social interaction. Moreover, TRAP-15q^+/-^ mice displayed impairments in their social preference index further substantiating their social deficits. This is notable since while TRAP-WT mice consistently demonstrated positive values indicative of a pronounced social preference, TRAP-15q^+/-^ mice exhibited a spectrum of responses, spanning from markedly negative to strongly positive values. Such responses may potentially be associated with the phenotypic behavioral variability characterizing human carriers [50]. Furthermore, the high variability in the penetrance of the microdeletion observed in humans [51], could contribute to the heterogeneity of social behaviors in the mouse model [49]. We also compared gender-specific alterations in social preferences and found that the effect of 15q13.3 microdeletion consistently affected both sexes and their impaired social behavior is conserved across groups.

After validating the behavioral impairments, we employed Psych-TRAP for whole brain automated alignment and quantification of cells activated after 3-CSI. We observed the activation mostly in cortical areas. Several regions are consistent with the activation patterns described in the literature, such as somatosensory areas (primary somatosensory area (SSp) and supplemental somatosensory area (SSs)) [26][27][28][29][30][52], which play a key role in sensory processing and experience-dependent behavioral adaptations [53][29]. The differences were not only limited to the overall area of SS, but also to their subregions, indicating layer-specific roles in sensory-motor integration [54][55].

Another key region which was strongly associated with social behavior and psychiatric phenotypes is anterior cingulate area (ACA), which has been validated consistently to play major roles in shaping social information [35][56] and serves as an information transfer region between limbic system and cognitive system [57]. Additionally, the higher activation in the RSP is natural due to receiving synaptic inputs from visual areas, hippocampus and thalamic nuclei and consequently playing a role in social behavior and in sensory and spatial information processing in psychiatric models [58][59].

It was surprising to observe higher activation in primary motor areas (Mop) in TRAP-15q^+/-^ mice, even though the movement and velocity across both groups remain unchanged. The role of MOp is to perform fine-motor controls and to coordinate inputs from and to other brain regions [60][61]. Higher activation of MOp might indicate potential involvement of the motor cortex in social behavior in psychiatric context. Another novel brain area that was activated in our experiments and was not directly linked to social behavior before is gustatory area (GU4), which plays role in taste encoding and decision making [62]. Gustatory areas are a part of associative areas that integrate multi-sensory information and act as a relay region [63]. Since TRAP-15q^+/-^ mice exhibited behavioral impairments and recruited several sensory cortical areas, gustatory areas may be involved in integrating visual and odor information, thereby showing higher activation patterns.

To reveal the subtypes of activated cells, we zoomed into SSp-tr and PFC regions, which were among the major regions exhibiting higher activation in the TRAP-15q^+/-^ mice. We first confirmed lack of activation for non-neuronal cells. We further determined subtypes of activated neurons, where we focused on GABAergic neuronal subtypes since their impairment has been known to be compromised in various psychiatric conditions modelled with genetic and environmental risk factors [16][33][64][65][66][67][15][23].

The role of PV+, SST+ and RELN+ neurons have been implicated in social behaviors [35][37][68][69][70][71][72], but their association with psychiatric-related impairments in 15q13.3 microdeletion has not been studied before. Interestingly, we observed a stronger activation of GABAergic component in the PFC after 3-CSI in the TRAP-15q^+/-^ mice, indicating a shift in E/I balance. This could be due to the diminished density of PV+ interneurons in the PFC, as they are major drivers in maintaining E/I balance [73]. Additionally, we revealed the profound increase in activation of SST+ and RELN+ neurons in SSp-tr and PFC brain regions, respectively. Surprisingly, PV+ neurons had much lower activation relative to SST+ neurons, in both TRAP-15q^+/-^ and the TRAP-WT mice, indicating larger role of SST+ neurons in this behavioral paradigm.

Limitations of Psych-TRAP approach includes using TRAP2 labelling, thus capturing Fos-based gene activity, making it harder to capture populations that express other IEGs, such as like *Arc* and *Npas4* [74]. Additionally, the concentration of the delivered 4-OHT, that binds to the iCRE and causes recombination, is important since it defines the total amount of labeled neurons. However, a minimum threshold of 4-OHT concentration needs to be achieved to observe positive signal while not causing unspecific labeling. We have optimized the concentration of 4-OHT in this study to 30 mg/kg to attain a detectable threshold. ABBA alignment also offers some limitations as it is hard to align and quantify sections that are broken, distorted or contain missing parts, since currently implemented automated counting marks it as ‘zero’ cells, if a brain area is not found.

Psych-TRAP enables users to label stimulus-specific populations in combination with mouse models of psychiatric risk factors. It offers whole brain labeling of stimulus-activated cells. These cell populations are labeled with a strong-temporal resolution in a permanent fashion and can be manipulated for functional studies in future, such as optogenetics and chemogenetics. To achieve spatial resolution, a user can also utilize Cre-Lox-mediated viral vectors in TRAP2 mice (without crossing with reporter mice), where the fluorescent labeling of active cells depends on the specific brain area into which the vector is transfected [75]. Psych-TRAP is a valuable and handy technique to address what neuronal types and where are activated by relevant clinical symptoms, discovering novel brain areas and molecular targets underlying complex psychiatric phenotypes. Overall, Psych-TRAP has strong potential to revolutionize modern labeling strategies and functional studies that utilize psychiatric models.

## Acknowledgements

The authors are grateful to the members of the Khodosevich groups for critical input during this project. We thank Dr. Katarina Dragicevic and Center for Health Data Science (HeaDS) for their help and guidance with the bioinformatic analyses. We wish to thank the BRIC Light Microscopy Core Facility for their constant support and excellent help during this study. We would like to thank Josefine Skov, Lilya Mork Jorgensen, Sarah Ahmed Al-Tewaj and Nicola Lanti for help with genotyping. This work was supported by the Novo Nordisk Foundation Hallas-Møller Investigator Grants - Emerging NNF16OC0019920 and Ascending NNF21OC0067146 to KK, and Lundbeck Foundation Collaborative Project R453-2024-507 to KK, DSFN 2022 Scholarstipendium to XS and Lundbeck Experiment Grant R480-2024-983 to MJ.

## Data availability

The datasets analysed for the current study are available from the corresponding authors on reasonable request.

## Author contributions

Research design and conceptualization, M.J. and K.K.; investigation, M.J.; methodology - behavior, M.J. and X.S.; methodology – immunohistochemical analyses, M.J., X.S., E.K.H., S.B.H.; methodology – ABBA alignment, M.J., E.K.H., discussion and troubleshooting, M.J., X.S., E.K.H., S.B.H.; funding acquisition, M.J., X.S., K.K.; writing - original draft, M.J.; writing - reviewing and editing, M.J., K.K.

---

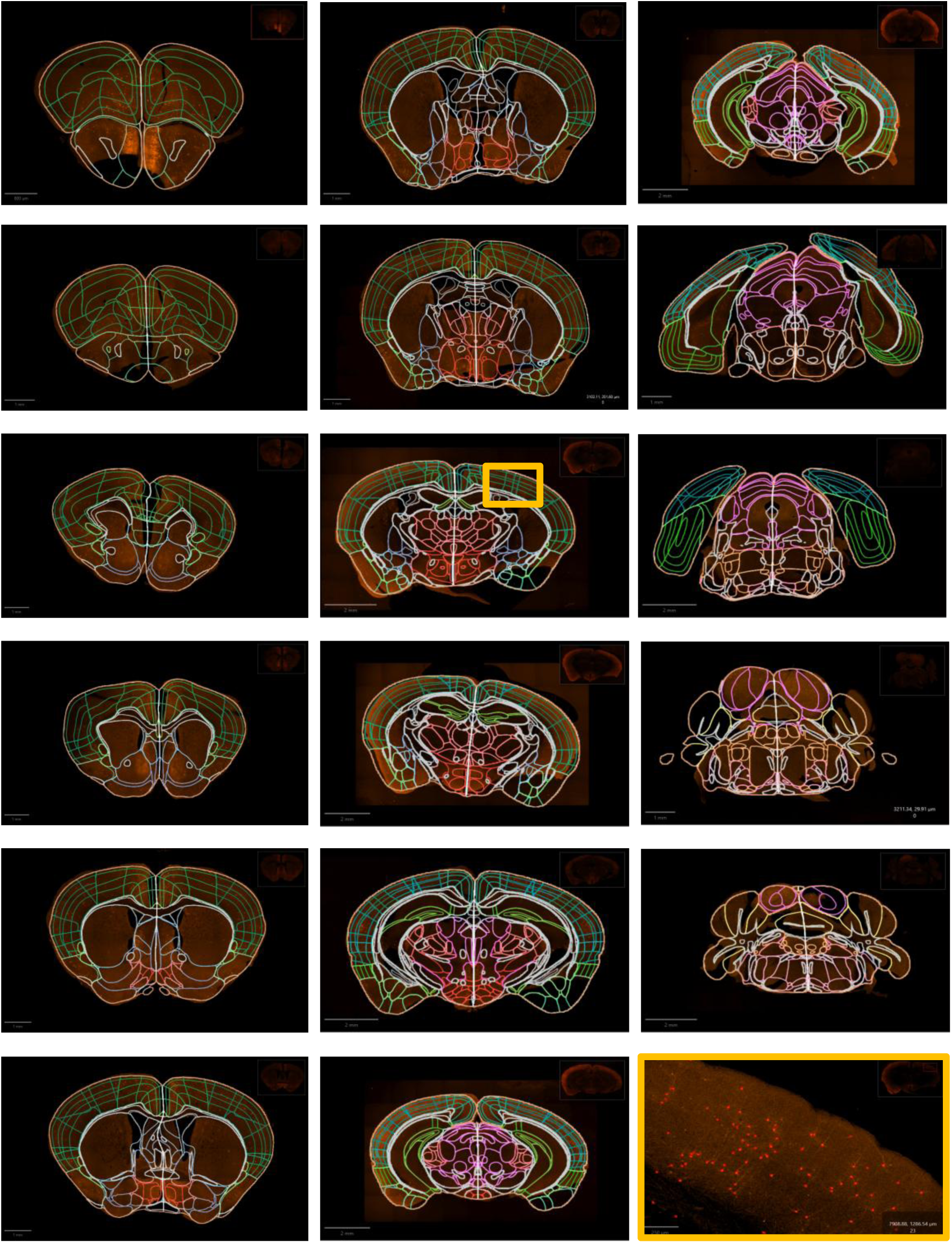

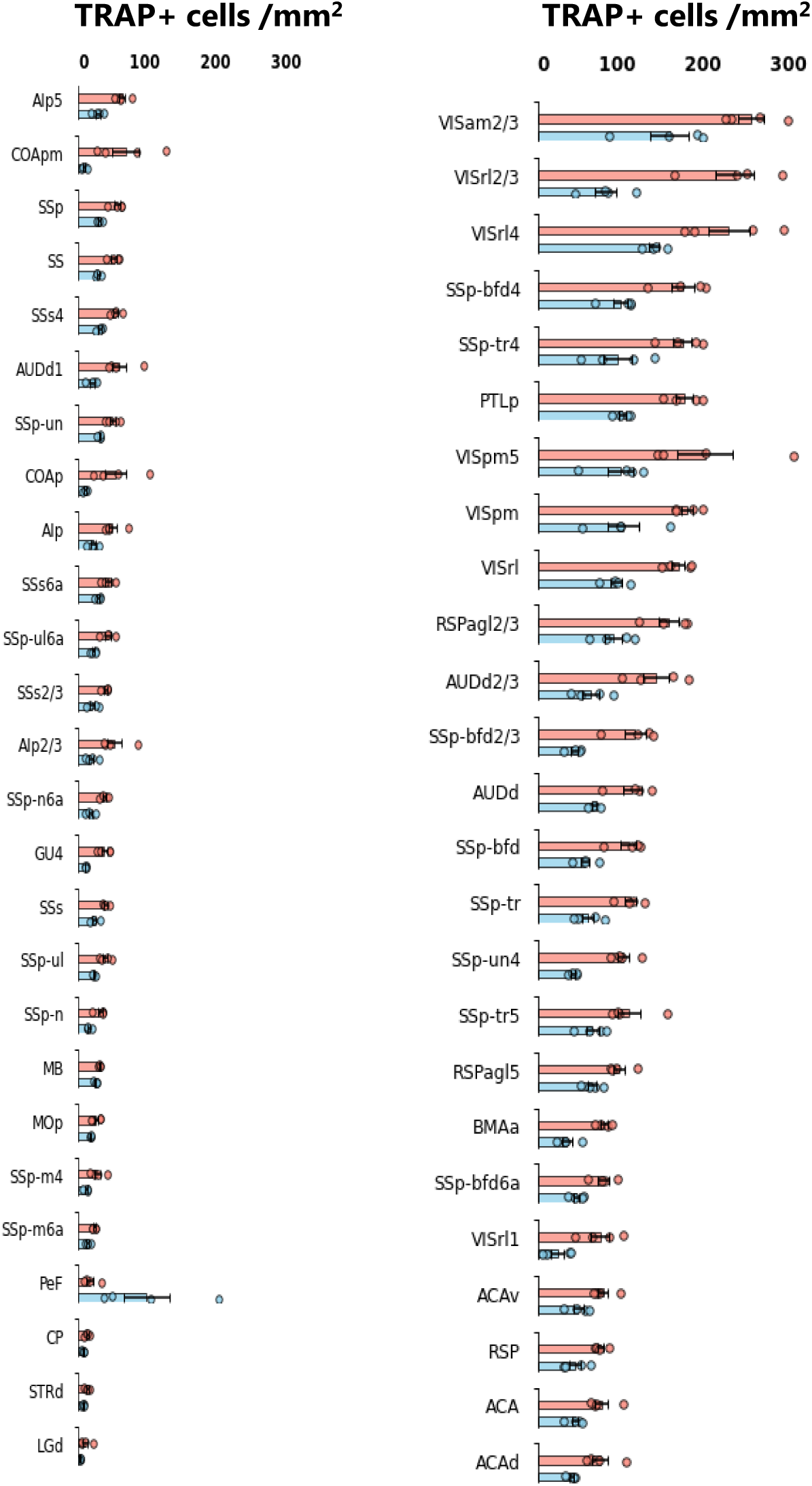

